# Inherited duplications of *PPP2R3B* promote naevi and melanoma via a novel *C21orf91*-driven proliferative phenotype

**DOI:** 10.1101/672576

**Authors:** Satyamaanasa Polubothu, Lara Al-Olabi, Daniël A Lionarons, Mark Harland, Anna C Thomas, Stuart Horswell, Lilian Hunt, Nathan Wlodarchak, Paula Aguilera, Sarah Brand, Dale Bryant, Philip Beales, Cristina Carrera, Hui Chen, Greg Elgar, Catherine A Harwood, Michael Howell, Dagan Jenkins, Lionel Larue, Sam Loughlin, Jeff MacDonald, Josep Malvehy, Sara Martin Barberan, Vanessa Martins da Silva, Miriam Molina, Deborah Morrogh, Dale Moulding, Jérémie Nsengimana, Alan Pittman, Juan-Anton Puig-Butillé, Kiran Parmar, Neil J Sebire, Stephen Scherer, Paulina Stadnik, Philip Stanier, Gemma Tell, Regula Waelchli, Mehdi Zarrei, Davide Zecchin, Susana Puig, Véronique Bataille, Yongna Xing, Eugene Healy, Gudrun E Moore, Wei-Li Di, Julia Newton-Bishop, Julian Downward, Veronica A Kinsler

## Abstract

The majority of the heredity of melanoma remains unexplained, however inherited copy number changes have not yet been systematically studied. The genetic environment is highly relevant to treatment stratification, and new gene discovery is therefore desirable. Using an unbiased whole genome screening approach for copy number we identify here a novel melanoma predisposing factor, familial duplications of gene *PPP2R3B*, encoding a regulatory unit of critical phosphatase PP2A. Significant correlation between expression of *PPP2R3B* in tumour tissue and survival in a large melanoma cohort was confirmed, and associated with a non-immunological expression profile. Mechanistically, construction and extensive characterization of a stable, inducible cellular model for *PPP2R3B* overexpression revealed induction of pigment cell switching towards proliferation and away from migration. Importantly, this was independent of the known microphthalmia-associated transcription factor *(MITF)*-controlled pigment cell phenotype switch, and was instead driven by uncharacterised gene *C21orf91*. Bioinformatic studies point to *C21orf91*as a novel target of *MITF,* and therefore a potential hub in the control of phenotype switching in melanoma. This study identifies novel germline copy number variants in *PPP2R3B* predisposing to melanocytic neoplasia, and uncovers a new potential therapeutic target *C21orf91* in the control of pigment cell proliferation.

## Introduction

Melanoma remains a major cause of morbidity and mortality. The majority of the heredity of melanoma remains unexplained, suggesting that multiple low prevalence variants are involved. Germline loss of function variants in *CDKN2A* in 2% of melanoma cases are the commonest known genetic predisposer^1–3^, with more recent discoveries of predisposing variants in genes *BAP1*^4^ and *POT1*^5^ at a similar frequency. With *CDKN2A*, affected individuals and their families are screened for melanoma under current guidelines^6^. Identification of new familial predisposing genes to melanocytic neoplasia is therefore desirable to identify families at higher risk of melanoma, however it also offers the potential to inform the understanding of the condition at molecular level. Whole genome germline copy number analysis has not yet been undertaken in melanoma cohorts. We sought to identify novel predisposing copy number changes for melanocytic neoplasia, using a rare disease approach.

Congenital melanocytic naevi (CMN; MIM 137550) is a rare congenital mosaic disorder of large and multiple moles, which predisposes affected individuals to melanoma. It is a valuable UV-independent model for the development of melanocytic naevi and melanoma, as the naevi develop *in utero*, and the melanoma most commonly arises post-natally but within the central nervous system^7^. It is also a good genetic model, as the causative somatic mutations are in classic melanoma hotspots in *NRAS* in 70%^8,9^, and *BRAF* in 7%^9,10^, with the remainder as yet unknown. Importantly for this study, despite the sporadic somatic nature of the disease, one third of cases have a first or second degree family history of CMN in the largest published cohort^11,12^, strongly suggesting predisposing germline genotypes to *NRAS*/*BRAF* somatic mutation in affected families. Previous work has delineated an enrichment of melanocortin-1-receptor (*MC1R*) variants in individuals with CMN^12^, again mirroring the genetic background of sporadic melanoma and underlining that this genetic model is UVR independent. We hypothesised that new predisposing copy number variants found via this rare disorder could predispose to melanoma in the normal population. Germline copy number in the CMN cohort was analysed using an unbiased whole genome approach, and findings validated in an independent adult sporadic melanoma cohort.

## Results

### Germline duplications involving *PPP2R3B* are found at increased frequency in individuals with melanocytic neoplasia

Using whole genome array comparative genomic hybridisation (CGH) of leukocyte DNA, duplications of Xpter were identified in 3/24 (12.5%) patients with CMN (**Figure 1a**), where only gene *PPP2R3B* was common to all three. No other novel copy number variant was seen in more than one patient. Control data from paediatric patients with other phenotypes from the same diagnostic testing facility identified duplications of this gene (with or without involvement of the two telomeric genes but not extending centromeric) in 13/4800, or 0.271%. Population data from normal individuals from the Database of Genomic Variants^13^ identified similar duplications in only 1/36,000, or 0.003%^14^, and 0.5% in nearly 7000 individuals in the MSSNG autism study^15,16^, with no difference between cases and controls (personal communication). Custom-designed TaqMan^®^ copy number assays (Thermo Fisher Scientific, USA) were insufficiently robust for validation due to highly repetitive DNA sequence in this area. Custom-designed multiple ligation-dependent probe amplification (MLPA, MRC Holland) and targeted next generation sequencing (NGS) of the four most telomeric genes on Xp confirmed the array findings, detecting an overall prevalence of duplications involving *PPP2R3B* but not downstream *SHOX* in 5% of a cohort of 125 individuals with CMN (**Figure 1c,d**). NGS of leukocyte DNA from an adult melanoma cohort identified the same germline duplications in 4/178 (2.2%) (**Figure 1b**), confirming this copy number variant as enriched in populations with melanocytic neoplasia. Regional similarity search across Xp22.33 revealed three segmental duplications 5’ of *PPP2R3B*, and a high density of SINE and LINE repeats, but no segmental duplications between *PPP2R3B* and *SHOX* (**Figure 1e**). Sanger sequencing of leukocyte DNA from 48 CMN patients and 48 controls did not detect any undescribed variants or haplotype differences (data not shown).

**Figure 1.**
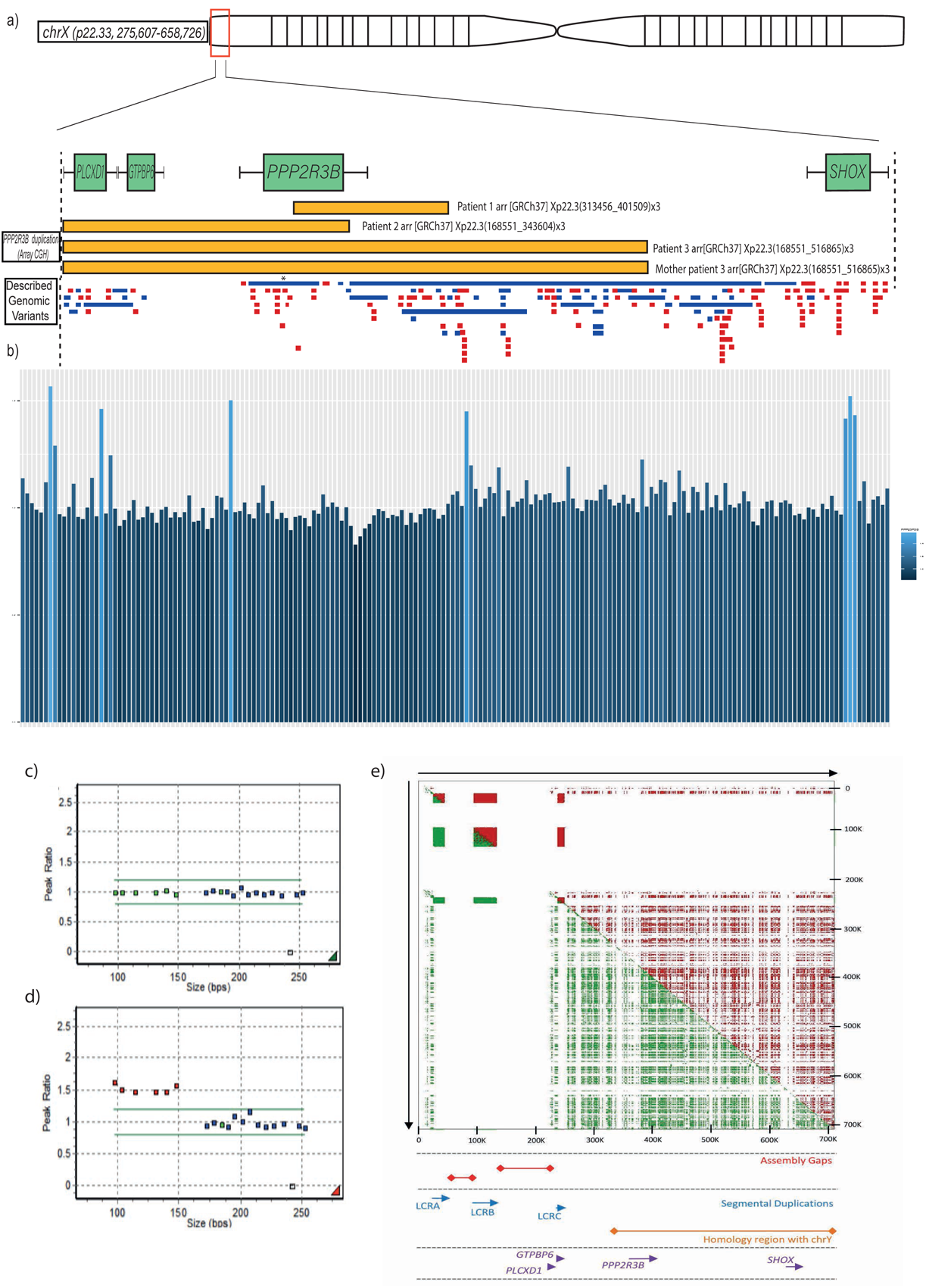
**Germline duplications involving *PPP2R3B* are found at increased frequency in individuals with melanocytic neoplasia** **(a)** Schematic of Xp22.33 region demonstrating the location of three novel duplications (yellow) found in 24 congenital melanocytic naevus (CMN) patients using whole genome array CGH of leukocyte DNA, with one identical parental duplication demonstrating inheritance. Described copy number variants from that region are shown below, duplications in blue, deletions in red, with each bar representing a single publication. The publication representing a duplication involving *PPP2R3B* described a single variant in a cohort of approximately 36,000 (asterisk, and see text for details), confirming that the CMN duplications are rare in the normal population. **(b)** *PPP2R3B* duplications in a UK non-syndromic melanoma cohort (n=179), and CMN cohort (n=8) shown by targeted next generation sequencing of *PPP2R3B*, in addition to the two telomeric genes (*GTPBP6 and PLCXD1*) and the next centromeric gene (*SHOX*). Data represent the ratio of corrected read depth (see text for details) across the whole of *PPP2R3B* with respect to the ratio across the whole of *SHOX.* Each bar represents an individual patient. *PPP2R3B* duplications called are shown in light blue: validation of the array CGH findings in the three CMN patients are clustered to the right of the figure, and new duplications in the melanoma cohort in the rest of the figure (n=4, 2.2%). **Validation of *PPP2R3B* duplications detected by array CGH** **(c-d)** Custom-designed MLPA^®^ ratio plots validating copy number measurement of *PPP2R3B* (4 probes) and the two telomeric genes *GTPBP6* and *PLCXD1* (one probe each), to the left of each figure and less than 150bp in length; control probes of greater than 160kb targeting genes of known normal copy number across the genome are shown to the right. Example of normal copy number for all genes **(c)**, and of a duplication of *PPP2R3B* and *GTPBP6* and *PLCXD1* in a CMN patient **(d)**. **Low-Copy Repeats at Xp22.33** **(e)** The upper panel depicts a regional similarity search across Xp22.33 with YASS software (http://bioinfo.cristal.univ-lille.fr/yass/index.php) both forwards (green) and backwards (red) revealing three segmental duplications (LCRA, LCRB and LCRC) 5’ of *PPP2R3B* and a high-density of SINE and LINE repeats. No segmental duplications are detected 3’ to *PPP2R3B* before *SHOX*. The assembly gaps (red), local genes (purple) and the homology region (orange) with the Y chromosome are indicated.

### Germline duplications of *PPP2R3B* lead to increased expression of PR70 in congenital melanocytic naevi

Owing to other potential genetic confounders within malignant tissue such as loss of Xp, effects of germline duplication on tissue expression *in vivo* was visualised in CMN tissue, known to have little somatic copy number variation^17^. Germline duplications were clearly associated with increased expression of PR70 in CMN tissue on immunohistochemistry compared to controls (**Figure 2a,b**).

**Figure 2.**
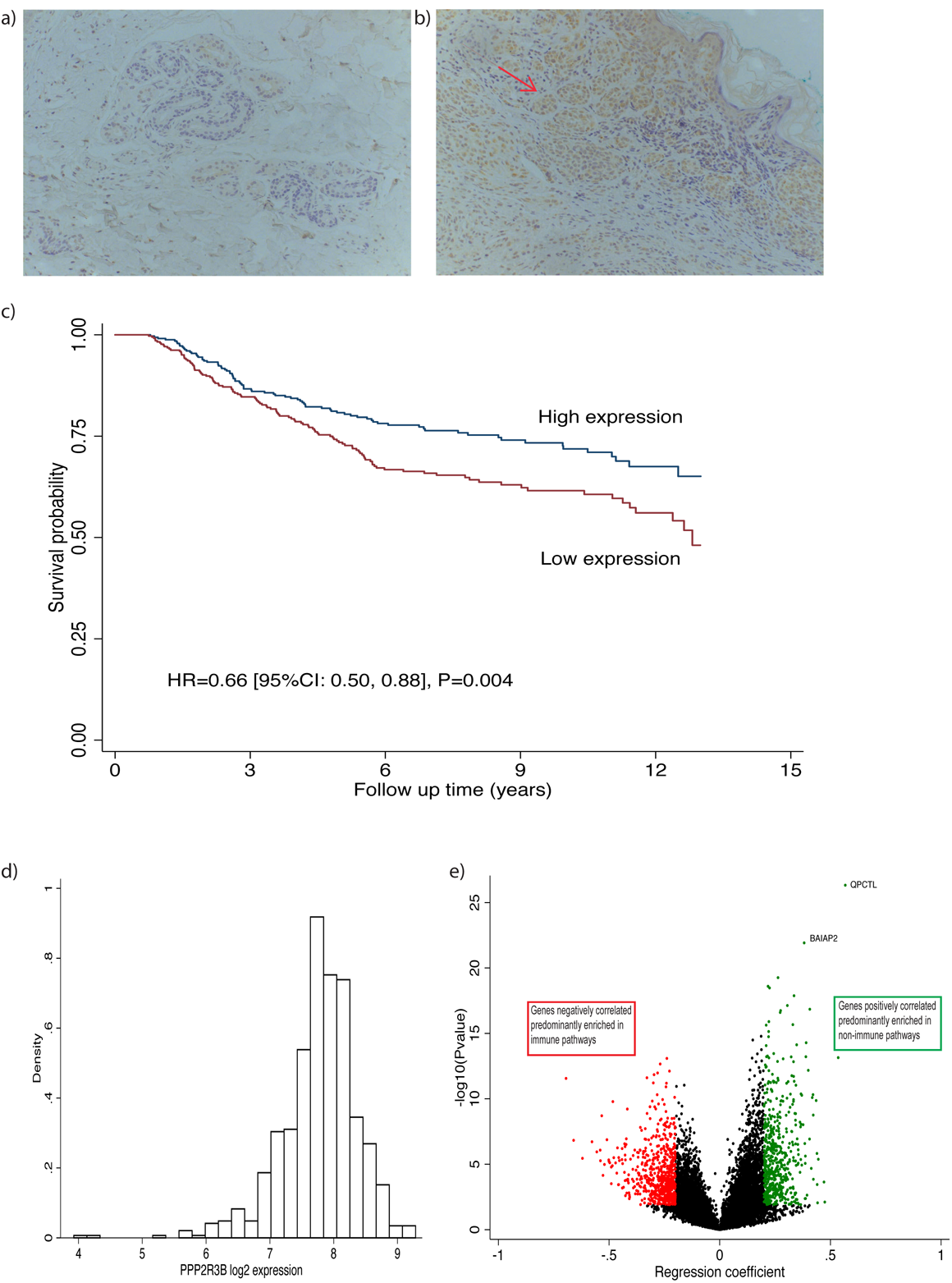
**(a-b) Germline duplications of *PPP2R3B* lead to increased protein expression within congenital melanocytic naevus tissue**, compared to that of normal copy number controls. Immunohistochemical staining of formalin fixed paraffin embedded CMN tissue demonstrates negative or very low intensity PR70 staining in a patient with confirmed normal copy number of *PPP2R3B* (a), and widespread moderate intensity staining throughout the cytoplasm of naevus cells in a patient with a confirmed germline *PPP2R3B* duplication **(b)**, both shown at 20X. Red arrow indicates nest of naevus cells with increased DAB staining. **(c-d) Increased *PPP2R3B* expression in melanoma tissue is correlated with improved melanoma-specific survival. (c)** Kaplan-Meier curve generated from transcriptomic data from 703 FFPE melanoma tumours from the Leeds Melanoma Cohort, hazard ratio (HR) =0.69, (95% confidence interval 0.50-0.88), p = 0.004. The effect remains significant after adjusting for age, sex, AJCC (American joint committee for cancer) stage, vascular invasion, site, *BRAF/NRAS* mutation status and tumour invading lymphocytes (TILs). **(d)** Log intensity distribution of *PPP2R3B* DASL probe (ILMN_1689720) is close to a normal distribution. **(e) Improved melanoma specific survival observed with increased expression of *PPP2R3B* appears not to be immune mediated.** Tumour expression of *PPP2R3B* correlates with expression of a large number of other genes in the genome: 596 positively correlated at FDR<0.05 with regression coefficient>0.20; 731 negatively correlated at FDR<0.05 with a regression coefficient<-0.2 **(e)**. The genes positively correlated with *PPP2R3B* are predominantly enriched in non-immune pathways, consistent with the lack of association between *PPP2R3B* expression and TILs or any specific immune cell score (Figure S1). The genes negatively correlated with *PPP2R3B* expression are predominantly enriched in immune pathways.

### Expression of PR70 is significantly associated with prolonged melanoma-specific survival (MSS)

Increased tissue expression of *PPP2R3B* in a larger melanoma cohort, although potentially influenced by various somatic changes in malignancy, was significantly associated with prolonged melanoma-specific survival (MSS) (**Figure 2c,d**). This effect remained significant after correction for known associations with survival, namely age, sex, AJCC (American joint committee for cancer) stage, vascular invasion, site, *BRAF/NRAS* mutation status and tumour invading lymphocytes (TILs). Unlike many known expression-survival associations in melanoma, transcriptomic pathway analysis demonstrated that this protective effect was not immune pathway-mediated (**Figure 2e, Tables S1,S2**), suggesting an alternative mechanism of action.

### Creation of a stable inducible overexpression system to study *PPP2R3B* overexpression

Detailed characterisation of the effects and mechanisms of *PPP2R3B* overexpression were modelled by creation of a stable inducible over-expression system in two melanoma cell lines SKMEL2 and SKMEL30 (**Figure 3a-c**). Induction robustly and reproducibly led to *PPP2R3B* mRNA overexpression, and protein product PR70 overexpression.

**Figure 3.**
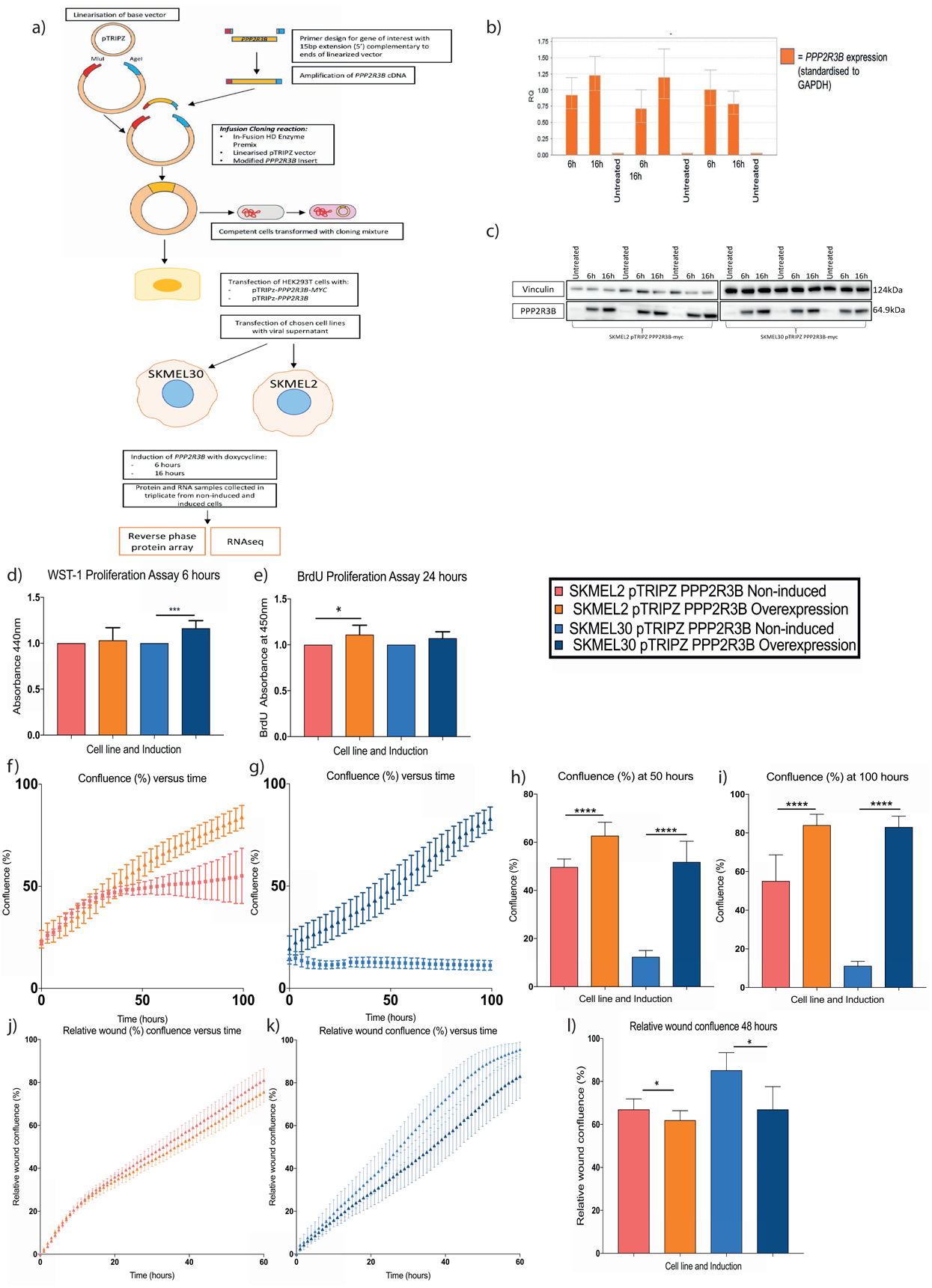
**(a-c) Generation of stable inducible overexpression model for *PPP2R3B* in melanoma cell lines SKMEL2 and SKMEL30.** Diagrammatic summary of stable inducible *PPP2R3B* overexpression system (SKMEL2 pTRIPZ *PPP2R3B* and SKMEL30 pTRIPZ *PPP2R3B)*, and generation of samples for reverse phase protein arrays (RPPA) and RNA sequencing **(a).** Validation of *PPP2R3B* overexpression in samples for Reverse Phase Protein Array and RNAseq, **(b)** qRT-PCR demonstrating increased *PPP2R3B* mRNA in induced cell lines at 4-6h and 16-18h hours (relative fold change in *PPP2R3B* expression, standardised to GAPDH, mean + SD of samples in quadruplicate), and, **(c)** Western blot confirming PR70 overexpression in induced cell lines at 4-6h and 16-18h hours with Vinculin loading control. **(d-j) Overexpression of *PPP2R3B* increases proliferation in melanoma cell lines.** Increased proliferation following *PPP2R3B* overexpression using WST1 proliferation assay at 6 hours **(d)**, and BrdU assay at 24 hours **(e)** (mean absorbance of colourmetric assay of eight replicates shown with error bar), and, by IncuCyte^®^ cell count proliferation assay over 100 hours, measuring confluence (%) versus time (hours) in SKMEL2 pTRIPZ *PPP2R3B* **(f)** and SKMEL30 pTRIPZ *PPP2R3B* **(g)** (mean confluence at each time point of eight replicates shown with error bars). The mean confluence in each cell line is shown at timepoints of 50h **(h)** and 100h **(i)**. **(j-l) Overexpression of *PPP2R3B* decreases cellular migration in melanoma cell lines.** Scratchwound assay in SKMEL2 pTRIPZ *PPP2R3B* **(j)** and SKMEL30 pTRIPZ *PPP2R3B* **(k)** leads to decreased relative wound confluence (%) versus time (hours) compared to non-induced controls. Mean relative wound confluence in each cell line shown at 48h **(l)** (mean relative wound confluence at each time point of eight replicates shown with error bars). Statistically significant values are indicated by a single asterisk (p<0.05), a double asterisk p<0.01, a triple asterisk (p<0.001) or a quadruple asterisk (p<0.0001). All p-values for Students t-test, Prism v7.0 (Graphpad).

### *PPP2R3B* overexpression leads to increased cellular proliferation and decreased migration in a 2D melanoma cell model

*PPP2R3B* overexpression led to clear pigment cell phenotype switching, most marked from 48 hours post induction onwards. This was measurable via significantly increased cellular proliferation by several alternative established methods (**Figure 3d-i**), and decreased migration in scratch assays couple with IncuCyte^®^ monitoring (**Figure 3j-l**).

### *PPP2R3B* overexpression does not significantly alter known melanoma signaling pathway activation

RNA sequencing pathway enrichment analysis identified suppression of the unfolded protein response and endoplasmic reticulum (ER) protein folding after induction of *PPP2R3B* (**Table S3**). Signaling pathway characterisation of 302 proteins using reverse phase protein arrays (RPPA, MD Anderson Core) pre- and post-induction of *PPP2R3B* demonstrated enrichment for mammalian target of rapamycin (mTOR) and hypoxia-induced factor 1 (HIF-1) signaling pathways, with prominent biological signatures of response to heat and stress (**Figure 4a-d, Table S4**). Functional protein association network analysis^18^ of differentially expressed proteins from RNA sequencing and RPPA (non-adjusted) created a strong network centred around these same pathways (**Figure 4l**). Immunoblotting provided validation of significantly decreased phosphorylation of AKT at 6-8 hours, however overall no dramatic effects on known melanoma signaling pathways were demonstrated (**Figure 4e-k**). Activation of CDC6, a known direct target of PR70^19^, was inconsistent across cell lines (**Figure 4k**). Increasing molar concentrations of PR70 up to a 1:1 ratio with that of the core PP2A enzyme increased PP2A activity towards its specific substrate pCDC6 in another cellular model, however further increases in concentration reduced phosphatase activity (**Figure S3**). PR70 was shown to be highly efficient at binding to the PP2A core enzyme when competing with another regulatory subunit B’γ1, and overexpression of V5-tagged PR70 in a mammalian cell line (C6) did not lead to free PR70 (**Figure S4**), suggesting either competitive binding of other PP2A holoenzymes or direct interactions with other effectors.

**Figure 4.**
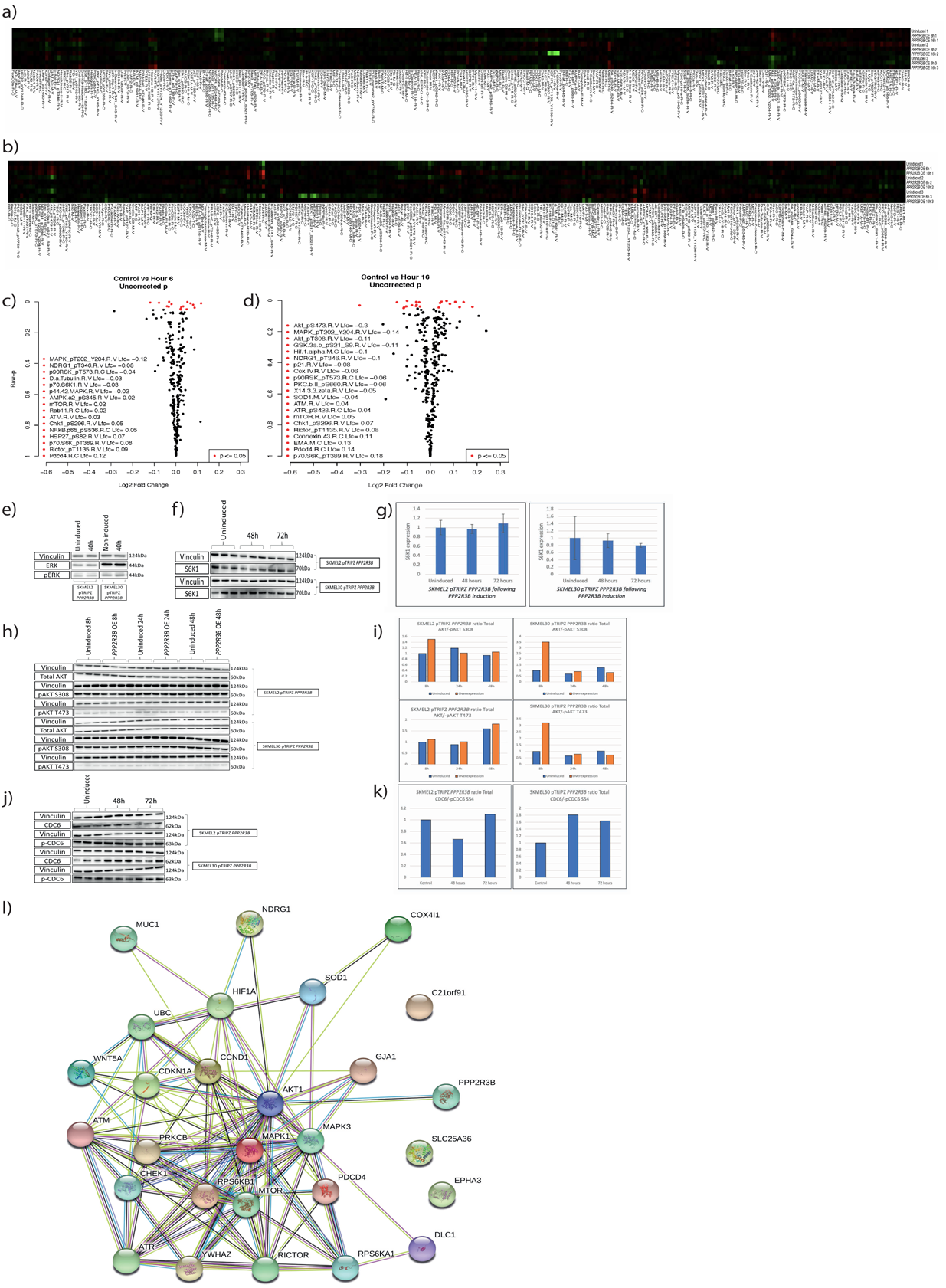
***PPP2R3B* overexpression affects mTOR/p70S6K1 and HIF-1 signalling pathways** **(a-b)** Heat map of protein expression observed by RPPA following overexpression of *PPP2R3B* in SKMEL2 pTRIPZ *PPP2R3B* **(a)** and SKMEL30 pTRIPZ *PPP2R3B* **(b)**, demonstrating low background activity as expected from a controlled cellular model. **(c-d)** Volcano plot of log fold change in protein expression versus p-value for differentially expressed proteins common to both cell lines following *PPP2R3B* overexpression. Unadjusted P-values < 0.05 are shown in red at 6h **(c)** and 16h **(d)**. **Overexpression of *PPP2R3B* dephosphorylates AKT, but does not consistently affect CDC6 activation.** Western blot of lysate from uninduced and induced SKMEL2 pTRIPZ *PPP2R3B* and SKMEL30 pTRIPZ cells, for ERK, phospho-ERK **(e)**, S6K1 and quantification **(f, g)**, AKT and phospho-AKT (S308 and T473) **(h)**, quantification of ratio of total to phospho-AKT **(i)**, demonstrates *PPP2R3B* overexpression decreases phosphyorylation of AKT at 6-8 hours, but does not lead to strong activation of known melanoma signalling pathways. Western blot of CDC6 and phospo-CDC6 **(j)** and quantification of ratio of total to phospho-CDC6 **(k)**, show *PPP2R3B* overexpression does not consistently alter CDC6 activation across the cell lines in this system, as confirmed by direct phosphatase activity measurements in another cellular model demonstrating that the interaction between PR70 and CDC6 may be dose-dependent **(Figure S3)**. **Strong network of known protein interactions implicated following *PPP2R3B* overexpression identifies *C21orf91* as a key new gene in these pathways.** Functional protein association network using STRING^15^ software demonstrating strong network of known signalling pathway interactions between all significant pre-adjustment RPPA hits and all significant RNAseq hits present in both cell lines following *PPP2R3B* overexpression at 16 hours **(l)**. *C21orf91* was the most differentially expressed gene in each cell line but is not currently known to be connected to these pathways.

### *PPP2R3B* overexpression leads to a significant and sustained rise in expression of the previously uncharacterized gene *C21orf91,* independent of *MITF*

Given the lack of clear signaling pathway activation in the presence of a clear proliferative phenotype, an alternative candidate mediator was sought by unbiased methods. In this regard, relatively unknown gene *C21orf91* (Refseq Gene ID:54149) was identified by RNA sequencing as the most significantly differentially-expressed gene in both cell lines, and validated at mRNA and protein levels (**Figure 5, S1, Table S5**). Crucially, knockdown of *C21orf91* by siRNA reversed the increased proliferation associated with induction of *PPP2R3B* expression (**Figure 6a-d**), firmly tying *C21orf91* to the pro-proliferative phenotype. Importantly, this increase was independent of microphthalmia-associated transcription factor (*MITF)*, master regulator of melanocyte transcription and phenotype switching (**Figure S2**), as witnessed by the lack of MITF overexpression at mRNA and protein levels upon induction of *PPP2R3B*.

**Figure 5.**
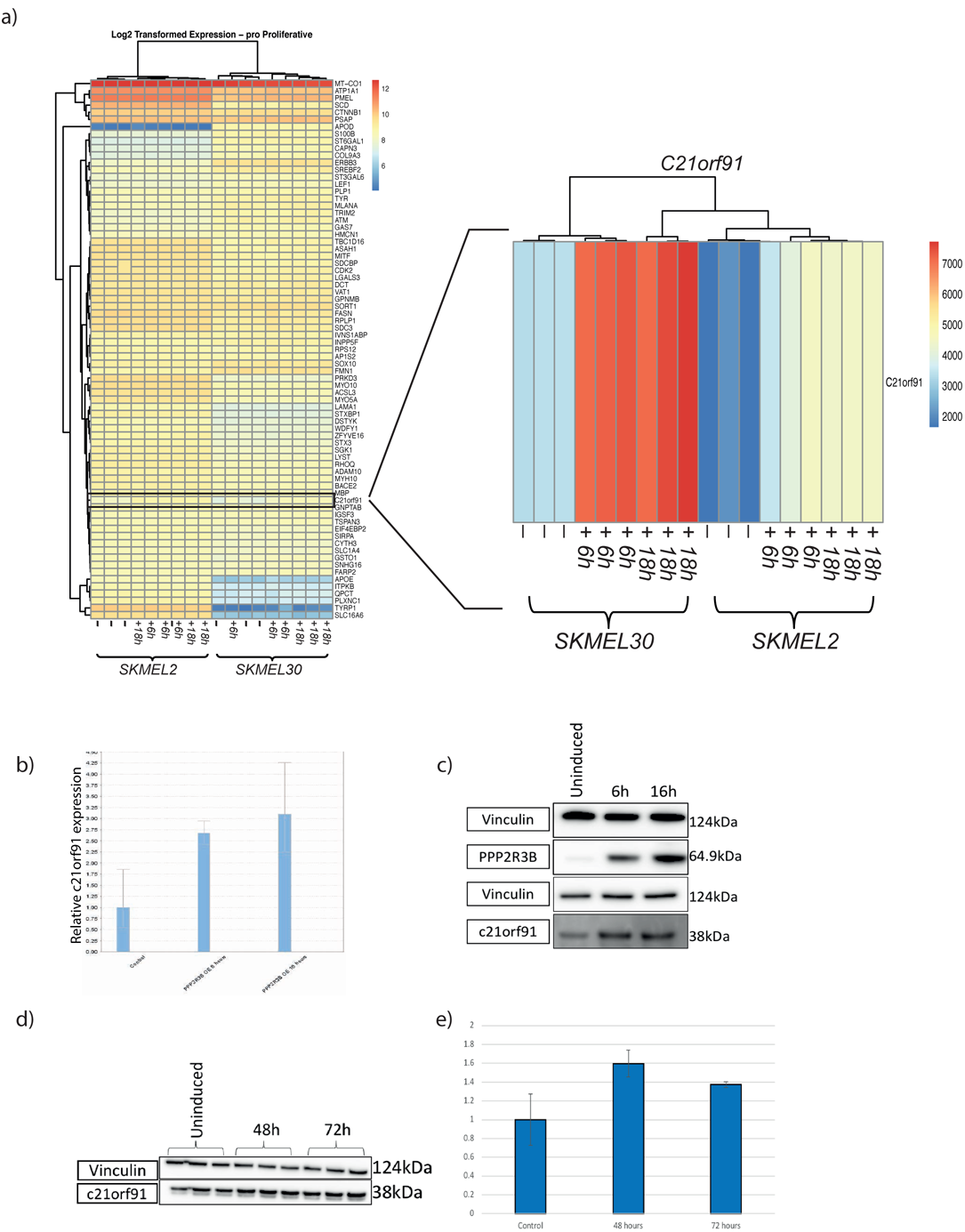
***PPP2R3B* overexpression leads to significant and sustained rise in expression of gene C21orf91.** *C21orf91* was the most differentially expressed gene on *PPP2R3B* induction common to both cell lines and at both time points by RNA sequencing (**Table S1**), other than *PPP2R3B* itself. (a) Heat map from pathway signature analysis of RNAseq data at 6-8h and 16-20h, focusing on pro-proliferative anti-invasive melanoma signature genes^38^, clearly demonstrating increased expression of *C21orf91* following *PPP2R3B* overexpression observed in both cell lines, at 6 and 16 hours. Validation of significantly increased *C21orf91* expression following *PPP2R3B* overexpression at 6h and 16h hours, shown by **(b)** qRT-PCR relative fold change standardised to *GAPDH*, mean + SD of samples in quadruplicate and **(c-d)** Western blots with loading control vinculin, and (d) quantification of fold change observed in *C21orf91* following overexpression of PR70.

**Figure 6.**
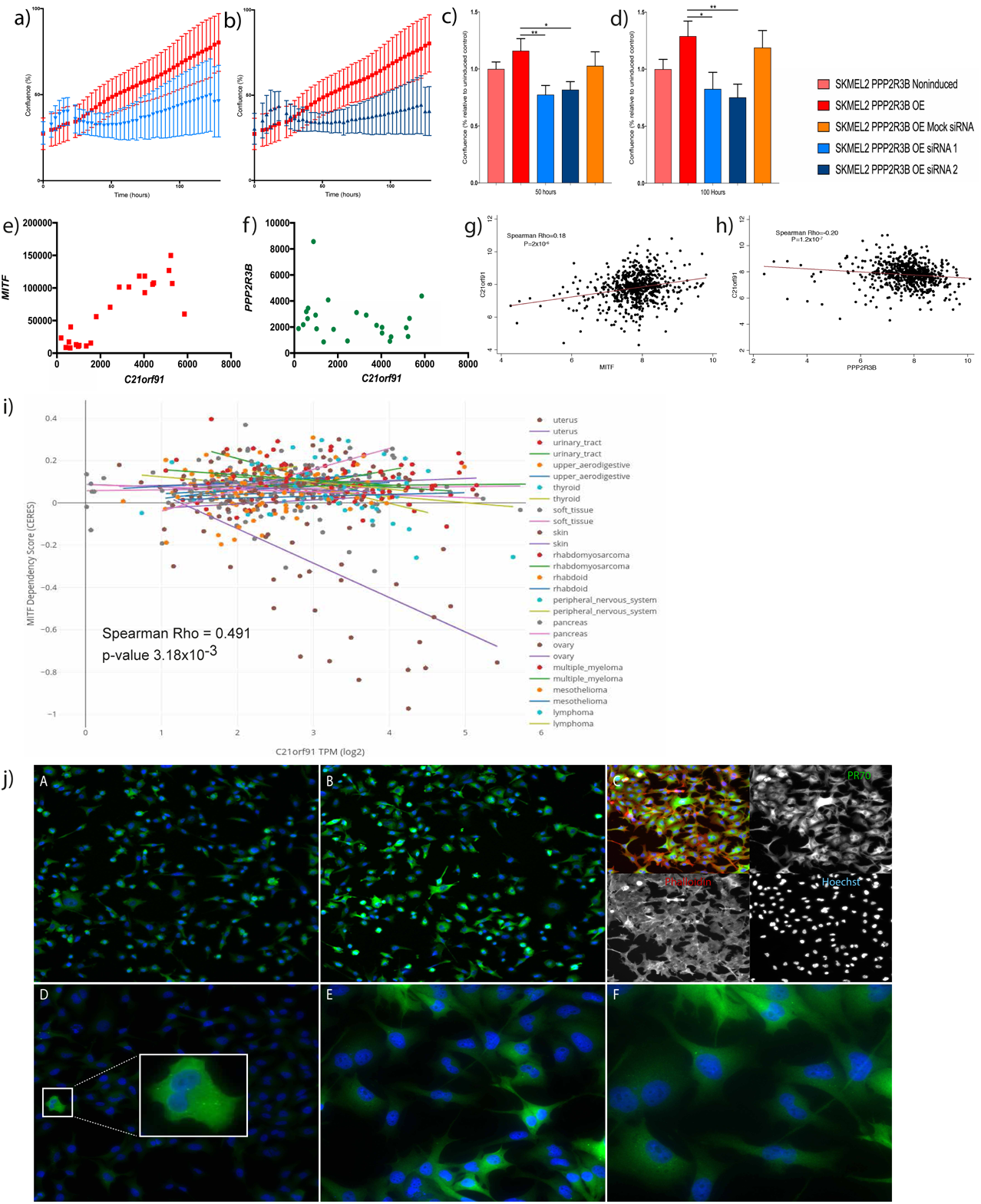
**Knockdown of *C21orf91* rescues increased proliferation associated with *PPP2R3* overexpression** IncuCyte^®^ proliferation assay confluence (%) versus time (hours) for *PPP2R3B*-induced SKMEL2 pTRIPZ *PPP2R3B,* with and without knock-down of *C21orf91*, by siRNA1 **(a)**, and siRNA2 **(b)**, with means taken at 50h **(c)**, and 100h **(d)** (mean confluence at each time point of eight replicates shown with error bar, standardised to uninduced control). Single asterisk p<0.05, double asterisk p<0.01 (Students t-test, Prism v7.0 (Graphpad)). ***C21orf91* and *MITF* expression are significantly positively correlated in melanoma**, independent of *PPP2R3B* Positive correlation demonstrated in independent transcriptomic data from a cohort of 24 human derived melanoma cell lines, Spearman Rho 0.18 (p = 2 × 10^−6^) **(e)** and in human melanoma tissue (Spearman Rho 0.8 (95% CI 0.6-0.9, p<0.0001) **(g)**. No correlation between expression of *C21orf91* and *PPP2R3B* in either cohort **(f)** and **(h)**, points to increased expression of C21orf91 as a new hub in melanocytic proliferation, not only a target of *PPP2R3B*. **Increased expression of C21orf91 is associated with genetic dependency on MITF in melanoma cell lines** CRISPR-Cas9 genome-scale knockout of *MITF* in melanoma cell lines (n=30) reveals increased expression of *C21orf91* in cells with greater MITF dependency **(i**), suggesting that *C21orf91* is downstream of *MITF*. Dependency score as described in https://depmap.org. **PR70 and C21orf91 are expressed throughout the cytoplasm with increased expression of C21orf91 in dividing cells** **(i)** Immunocytochemistry of SKMEL30 pTRIPZ *PPP2R3B* in uninduced cells at 10X **(A)** and induced cells at 10X **(B)** and 20X **(C)** confirms increased PR70 and C21orf91 expression throughout the cytoplasm following induction of *PPP2R3B*. PR70 is stained with Alex Fluor^®^ 488 (green) secondary antibody, and nuclei stained with Hoescht (blue) and in (C) Phalloidin (actin) is visualised with a conjugated Alex Fluor^®^ 647 (far red) antibody. C21orf91 expression is increased in dividing cells, 10X **(D)**, 20X **(E)**, 40X **(F)**, stained with Alex Fluor^®^ 488 (green) secondary antibody, and Hoescht nuclear stain (blue).

### *C21orf91* expression is positively correlated with *MITF* expression in melanoma

*Given that MITF is the known master regulator of pro-proliferative phenotype switching in melanoma, and given that C21orf91 was not operating via MITF, we considered whether MITF could instead be operating via C21orf91. C21orf91* and *MITF* expression were indeed found to be significantly positively correlated in independent transcriptomic datasets from both melanoma cell lines and the melanoma patient cohort (**Figure e-h**), implying at least a key role for *C21orf91* in the pro-proliferative state of melanoma, and potentially that *MITF* can operate via *C21orf91*. *MITF* dependency score and *C21orf91* expression in melanoma cell lines were also found to be significantly associated in melanoma cell lines **(i)**, data extrapolated from the Cancer dependency Map (Broad Institute) ^20–23^. PR70 subcellular localization by confocal microscopy was pan-cytoplasmic, including but not restricted to endoplasmic reticulum as previously suggested^24^, and increased but unaltered in distribution by overexpression (**Figure 6i).** C21orf91 expression was also demonstrated throughout the cytoplasm, and both PR70 and C21orf91 were noticeably increased in dividing cells (**Figure 6i**).

## Discussion

Copy number in the genome has been less systematically explored than sequence variation due to technical constraints^25^, with copy number variation enriched in areas of low genome mappability^26^, however copy number variants (CNVs) are prevalent in genes for cell communication and RAS-pathway signaling, including serine threonine kinases and phosphatases^27^. Using a rare disease cohort, we identify here new germline duplications in the pseudoautosomal region 1 of the X chromosome which predispose to melanocytic neoplasia. The minimal common area included only gene *PPP2R3B,* which encodes PR70, ubiquitously expressed in the cytoplasm, and one of the ß” family of regulatory units of the critical phosphatase and regulator of the cell cycle PP2A^28,29^. PP2A is a heterotrimeric holoenzyme consisting of a structural A subunit, a catalytic C subunit, and a regulatory B subunit^28,30,31^. The numerous non-homologous regulatory subunits are classified into B, B’, B’’ and B’’’ subfamilies, implicated in control of enzyme activity and substrate specificity^28,32^. PP2A operates via key effector pathways RAS/MAPK, Wnt and AKT/mTOR^28,29^. As such, PP2A activity is intimately involved in malignancy and response to treatments^33^, and is a major focus of potential therapeutics^33–36^.

Our data demonstrate that *PPP2R3B* overexpression promotes proliferation of melanocytes, which could explain the predisposition to the development of a clinically-apparent melanocytic naevus or melanoma in the context of a somatic mutation in a melanocyte. Alternatively, the pro-proliferative germline environment could itself predispose to somatic mutation in the skin, via increased cell division or alteration of cell cycle regulation and the associated effects on DNA repair. Interestingly, in parallel, our data demonstrate clearly that increased *PPP2R3B* expression simultaneously improves survival in melanoma, with a similar significant protective effect of *PPP2R3B* expression seen in urothelial cancer and pancreatic cancer datasets from the TCGA database. In the melanoma cohort studied here, this protection is mediated via a non-immunological mechanism, and could potentially operate via phenotype switching towards proliferation and away from migratory potential. In support of this data, a comprehensive study of the role of *PPP2R3B* in melanoma at tumour as opposed to germline level found the region to be copy-number sensitive, with loss of the inactivated X in females and decreased expression in males linked to decreased distant metastasis-free survival. The authors proposed that the copy number sensitivity of this locus could explain the gender differences in melanoma incidence and survival^37^.

Pigment cell phenotype switching is classically controlled by a reciprocal relationship between *MITF* and *POU3F2*^38–40^, however here was MITF-independent, and appears instead to be driven by the relatively unknown gene *C21orf91.* Although this mechanism could involve the AKT/mTOR/pS6K pathway, signalling pathway alterations were largely unimpressive, and indeed protein modelling demonstrated a decrease in phosphatase activity with an increasing ratio of PR70 to the core enzyme. Furthermore, significant association between *MITF* and *C21orf91* expression in a melanoma cohort and pooled melanoma cell lines offers the intriguing possibility that *MITF* may itself operate via *C21orf91*, identifying this as a potential new hub in control of melanoma cell cycle regulation. Supportive evidence for a role of *C21orf91* in this field includes its previous identification within a pro-proliferative anti-invasive transcriptomic signature in melanoma^41^, a central role in cell phenotype determination in neurological development^42^, and recognition as one of 180 key molecules in cross-species protein networks around the Ras-MAPK/PI3K pathways^43^.

Due to the highly repetitive nature of this region of the genome near the telomeric end of Xp, we found targeted NGS to be the most reliable way to detect and confirm duplications of *PPP2R3B*. Future screening of larger melanoma cohorts and families will likely require development of a diagnostic-grade test from the point of view of cost, which will allow assessment of the frequency across different cohorts, and of the penetrance of the melanoma phenotype associated with this variant.

## Conclusions

We identify here germline duplications in the gene *PPP2R3B* predisposing to naevogenesis and melanoma in an important proportion of cases. Duplications increase melanocytic tissue expression of the protein product PR70, which confers a survival advantage via promotion of a pro-proliferative and anti-migratory pigment cell phenotype, itself driven by a novel MITF-independent mechanism mediated by *C21orf91*. This work offers novel insights into both the origins and behaviour of melanocytic neoplasia, and identifies *C21orf91* as an important new and potentially targetable fulcrum in the control of proliferation.

## Methods

### Patient recruitment and sample collection

Congenital melanocytic naevi recruited from the department of paediatric dermatology, Great Ormond Street Hospital, UK and melanoma cohort from Professor Julia Newton-Bishop, University of Leeds. DNA was extracted from blood samples using standard methods in all cohorts.

### Ethical Approval

All participants gave written informed consent as part of ethically approved studies

The study of CMN genetics was approved by the London Bloomsbury Research Ethics Committee (REC) of Great Ormond Street Hospital (GOSH)/UCL Institute of Child Health (ICH). Samples from the Leeds Melanoma cohort was approved by the North East – York Research ethics committee (Jarrow, Tyne and Wear, UK).

### Array CGH

Whole genome array comparative genomic hybridization (CGH) was performed as per the manufacturer’s instructions on 29 germline DNA samples from the CMN cohort, using Roche Nimblegen 135K oligonucleotide arrays and sex-matched commercial pooled controls. 1-3 μg of patient DNA and control DNA (Megapool reference DNA, Male – EA-100M, Female – EA-100F, Kreatech, The Nederlands) was labelled with fluorescent dyes Cy3 (pink) and Cy5 (blue) respectively using NimbleGen Dual-Color DNA Labeling Kit according to the manufacturer’s protocol. Labelled samples were hybridized to the oligonucleotide array using the NimbleGen Hybridization System. Two-color array scanning was performed using a Molecular Devices GenePix 4400A (Molecular Devices inc, Sunneyville, CA, USA) at a resolution of 2.5 microns, with comparison of fluorescence from both dyes. Deva software from NimbleGen was used for data extraction, and data analysed using InfoQuant CGHFusion (version 5.7.0 on initial samples and later 6.1.0) or later samples Chromosome Analysis Suite 4.0 (ChAS 4.0, Thermofisher Scientific). Array CGH data has been submitted to http://www.ncbi.nlm.nih.gov/clinvar/.

### MLPA^®^, DNA sample preparation and internal controls

MLPA^®^ was performed as per MRC-Holland MLPA^®^ DNA protocol version MDP-v003, optimized using 25ng of DNA per well, and a 1 in 2 dilution of the customised probemix, using four probes for *PPP2R3B,* and one each for *GTPBP6* and *PLCXD1* designed to dovetail with the probes in the reference SALSA^®^ MLPA^®^ probemix P200-A1 (MRC-Holland). Capillary electrophoresis of diluted PCR products with ROX-500 size standard was performed on an ABI 3730xl, and data analysed using GeneMarker^®^ v2.4 (SoftGenetics, USA). Each plate contained the same controls of normal copy number and duplication for standardisation.

### Targeted Next Generation Sequencing Panel

A SureSelect targeted panel (Agilent Technologies, UK) was designed to capture the whole genomic region encompassing the last 4 genes on the pseudoautosomal region of X and Y chromosomes, chrX: 198061-607558 chrY: 148061-557558 (hg19, GRCh37). Library preparation was carried out using the SureSelectXT kit under manufacturers instructions (Agilent Technologies, UK). Samples were run on a NextSeq instrument 500/550, read length of 2×150 bp (Illumina, USA). In total 183 samples were processed including a melanoma cohort (n= 168), CMN cohort (n=5) and control samples from the ALSPAC study (n=3). BAM files were inputted to DeepTools MultiBamSummary using the PPP2R3B_Moderately_Stringent_1_covered.bed probe coordinates file. Coverage across probe regions within the gene hg19 coordinates were extracted and averaged. 3 Control samples were used to create ‘normal expected’ coverage ratios of the genes compared to *SHOX*. All samples were normalised compared to these ratios and R studio was used to visualise and calculated gene coverage data across all samples.

### Immunohistochemistry

The expression of protein product PR70 was assessed in formalin fixed paraffin embedded naevus tissue from patients with known *PPP2R3B* copy number status, both duplications and normal copy number, using an antibody against PR70 (see Table 1 for source, dilution) on the automated LEICA-Bond-Max immunostainer.

**Table 1.** Antibodies

| **Antibody** | **Source** | **Catalogue Number** | **Dilution** |
| --- | --- | --- | --- |
| **PPP2R3B** | Rabbit | Abcam, ab72027 | 1:1000 Western Blot<br>1:500<br>Immunocytochemistry |
| **MITF** | Mouse | Abcam ab3201 | 1:1000 |
| **Vinculin** | Mouse | Cell signaling<br>technology #4650S | 1:1000 |
| **C21orf91** | Rabbit | Altas Antibodies,<br>HPA018288 | 1:1000 Western Blot<br>1:500<br>Immunocytochemistry |
| **MYC-tag** | Mouse | Abcam ab32 | 1:2000 Western Blot<br>1:500<br>Immunocytochemistry |
| **S6K1** | Rabbit | Abcam ab32529 | 1:1000 |
| **AKT** | Rabbit | Cell signaling<br>technology #4685 | 1:1000 |
| **Phospho AKT S473** | Rabbit | Cell signaling<br>technology #9271 | 1:1000 |
| **Phospho AKT T308** | Rabbit | Cell signaling<br>technology #13038 | 1:1000 |
| **MYC** | Mouse | Abcam ab32072 | 1:1000 |

### Cell lines and culture

SKMEL2 and SKMEL30 melanoma cell lines were obtained from Cell Services, Francis Crick Institute. SKMEL2 were cultured in DMEM 10% FBS Penicillin streptomycin and glutamine cells incubated at 37°C 10% CO2. SKMEL30 were cultured in RPMI 10% FBS Penicillin streptomycin and glutamine cells incubated at 37 50% CO2. Normal *PPP2R3B* copy number of both cell lines was verified as described above.

### Generation of stable inducible *PPP2R3B* cells lines

*Human myc-FLAG tagged PPP2R3B ORF clone from Origene* (RC222908) was linearised and the insert DNA amplified using modified primers generating an N-terminal Myc tag. The In-Fusion^®^ HD Cloning system (Takara, 638909) was used to allow directional cloning of the *PPP2R3B* insert into the AgeI-MluI site of the lentiviral vector pTRIPZ (GE Healthcare) resulting in the final *PPP2R3B* (tet-ON) construct, without the TurboRFP or shRNAmir-related elements of the parental pTRIPZ plasmid. Transduction of HEK 293T cells with pTRIPZ-*PPP2R3B*, psPAX2 and pMD2.G plasmids using Lipofectamine 2000 generated lentiviral particles used to infect SKMEL2 or SKMEL30 target cells using polybrene. Stable cell lines were selected using puromycin.

### Sample preparation for Reverse Phase Protein Array and RNAseq

SKMEL30 pTRIPZ-PPP2R3B and SKMEL2 pTRIPZ-PPP2R3B cells were seeded and treated with doxycycline 1ug/ml for either 6 hours or 16 hours alongside untreated controls. Samples were collected in triplicate for both RNA extraction and protein lysis. RNA extraction and protein lysis was performed as described below. Confirmation of *PPP2R3B* induction in all replicates was performed by qPCR of reverse transcribed cDNA from extracted RNA, and Western Blotting for PPP2R3B.

### Reverse Phase Protein Array – MD Anderson

Protein samples were diluted to 1.5ug/ul with complete lysis buffer. Samples were submitted to MD Anderson, Core Facility. Reported intensity values were log transformed to approximate normality and comparisons were performed using an un-paired t-test.

### RNAseq

RNA integrity calculated using a Bioanalyser (Agilent). Library preparation using KAPA mRNA HyperPrep Kit (Roche) was automated using the Hamilton robot. Libraries were sequenced using a NextSeq 500 (Illumina, San Diego, US) with a 43-bp paired-end run. Data were trimmed for 3’ adapter sequences using Cutadapt 1.9.1, after which they were aligned to the ensemble GRCh38 release 86 human transcriptome using STAR 2.5.2a. Individual lane level replicates were merged using Samtools 1.8, after which raw gene counts were estimated using RSEM 1.3.0 and normalisation and differential expression calling was then performed using DESeq2. A corrected p-value of let than or equal to 0.05 was deemed to be significant. Pathway analyses based on genes reported in the various analyses were performed using Metacore (Clarivate Analytics)

### Quantitative Real Time PCR

RNA was extracted using RNeasy mini kit (Qiagen, 74104), as per manufacturer’s instructions. 1ug of total RNA was reverse transcribed to cDNA using Quantitect reverse transcription kit (Qiagen, 205311), and quantitative real time PCR was performed, using Taqman probes and the StepOnePlus Real-Time PCR System (ABI, Abilene, TX, USA) (see Table 2 for details of Taqman probes). Relative fold change to the housekeeping gene *GAPDH* was calculated by ΔΔCT method.

**Table 2.** Taqman^®^ probes

| **TaqMan® Probe 20X** | **Source, Catalogue Number** |
| --- | --- |
| ***PPP2R3B*** | Hs00203045_m1 |
| ***C21orf91*** | Hs00213743_m1 |
| ***GAPDH*** | Hs02786624_g1 |
| ***MITF*** | Hs01117294_m1 |

### Western Blotting

Protein lysate was quantified using Bradford protein assay, and 20ug of each sample was run on a 10% Bis-Tris protein gel. PVDF membrane was activated in methanol prior to transfer using a wet transfer reservoir. The membrane was washed three times in Tris Buffered Saline-Tween (TBST), then blocked using 3% albumin at room temperature for one hour. Membranes were then incubated in primary antibody in 3% albumin overnight at 4°C (see Table 1 for source, dilution). Membranes were washed three times in TBST then incubated in corresponding HRP-tagged secondary antibody in 5% milk for 1 hour at room temperature. Membranes were washed a further three times in TBST before adding ECL for two minutes followed by measurement of chemiluminescence using Amersham Imager 600. Intensity of bands was measured and quantified using Image J software.

### Immunocytochemistry

Cells were seeded at a density of 4 × 10^4^ cells per well in laminin coated 4 well chamber slides (Millicell^®^ EZ slide, Merck Millipore, PEZGS0816). Cells were washed in cold PBS and fixed using 4% paraformaldehyde (PFA), followed by permeabilisation using 0.1% Triton X100. Slides were washed three times with PBS and incubated with blocking solution (1% Bovine serum albumin, 10% Donkey serum, 0.3M Glycine, 0.1% PBS Tween) for 1 hour at room temperature followed by primary antibody in blocking solution overnight at 4 degrees (see Table 1 for source, dilution). Slides were washed once with 1:10,000 Hoechst for nuclear staining, then washed three times with 0.1% PBS Tween before incubation with a fluorescent tagged appropriate secondary antibody at room temperature for one hour in the dark. Slides were again washed three times with 0.1% PBS Tween and were mounted prior to analysis using a Leica upright fluorescent microscope.

### Proliferation Assays

#### WST1 proliferation Assay

SKMEL2 pTRIPZ *PPP2R3B* and SKMEL30 pTRIPZ *PPP2R3B* cells were seeded into a 96 well plate at a density of 1 × 10^4^ cells per well. *PPP2R3B* overexpression was induced using doxycycline (1ug/ml) alongside un-induced controls. Plates were incubated for 48 hours at 37°C, prior to the addition of WST-1 reagent. Plates incubated in the dark at 37 °C for a further two hours. Absorbance of plates was read in a spectrophotometer at 450nm and 620nm. Results were adjusted for absorbance of blank media, and absorbance of WST1 dye (620nm) and averaged between the twelve wells for each condition. Absorbance was averaged between 12 replicates for each condition (mean + SD) and statistical significance was calculated by Students t-test.

#### BrDU Proliferation Assay

BrDU assay was carried out using BrdU Cell Proliferation ELISA Kit, (Abcam, ab126556), as per the manufacturer’s instructions. SKMEL2 pTRIPZ *PPP2R3B* and SKMEL30 pTRIPZ *PPP2R3B* cells were seeded into a 96 well plate at a density of 2 × 10^5^ cells per well. *PPP2R3B* overexpression was induced using doxycycline (1ug/ml) alongside un-induced controls, and suggested assay controls. BrdU reagent diluted in media was added to all wells and plates were incubated for 24 hours at 37°C, at which point media was removed and Fixing Solution added to each well. Plates were incubated at room temperature for 30 minutes, then blotted dry before washing three times with Wash Buffer. Anti-BrdU monoclonal Detector Antibody was each well and plates incubated for 1 hour at room temperature. Plates were washed a further 3 times with Wash Buffer before adding peroxidase Goat Anti-Mouse IgG Conjugate and incubating for 30 minutes at room temperature. Plates were washed a final three times with Wash Buffer and once with distilled water before the addition of TMB Peroxidase Substrate Pipette. Plates were incubated for 30 minutes at room temperature in the dark. Stop Solution was added to each well and plates were absorbance at 450 nm was measured using a spectrophotometer. Absorbance was averaged between 12 replicates for each condition (mean + SD) and statistical significance was calculated by Students t-test.

#### IncuCyte^®^ Cell Count Proliferation Assay

SKMEL2 pTRIPZ *PPP2R3B* and SKMEL30 pTRIPZ *PPP2R3B* cells were seeded into a 96 well ImageLock plate at a density of 1 × 10^4^ cells per well. *PPP2R3B* overexpression was induced using doxycycline (1ug/ml) alongside un-induced controls, a total of 12 replicates per condition. The plate was then analysed using the IncuCyte**^®^** live-cell analysis system and set up to acquire 10X phase contrast images at a scanning interval of 60 minutes for 5 days, measuring percentage confluence. Confluence was averaged between 12 replicates for each condition (mean + SD) and statistical significance was calculated by Students t-test.

### *C21orf91* Knockdown

SKMEL2 pTRIPZ *PPP2R3B* were seeded into a 96 well ImageLock plate at a density of 1 × 10^4^ cells per well. *PPP2R3B* overexpression was induced using doxycycline (1ug/ml) alongside un-induced controls, a total of 12 replicates per condition. Selected cells were transfected with Lipofectamine™ RNAiMAX using two different *C21orf91* siRNAs at a concentration on 10nm or a Scrambled siRNA (Origene, SR310041). The plate was then analysed using the IncuCyte^®^ live-cell analysis system and set up to acquire 10X phase contrast images at a scanning interval of 60 minutes for 5 days, measuring percentage confluence. Confluence was averaged between 12 replicates for each condition (mean + SD) and statistical significance was calculated by Students t-test.

### Migration Studies

#### Scratch Wound Assay

SKMEL2 pTRIPZ *PPP2R3B* and SKMEL30 pTRIPZ *PPP2R3B* cells were seeded into a 96 well ImageLock plate at a density of 1 × 10^5^ cells per well. *PPP2R3B* overexpression was indut.ced using doxycycline (1ug/ml) alongside un-induced controls, 12 replicates per condition. Plates were incubated at 37 degrees until all wells were confluent. WoundMaker™ was used to create a scratch in each well as per manufacturer’s instructions. Plates were washed with PBS to remove any debris and fresh media added to each well. The plate was then analysed using the IncuCyte**^®^** live-cell analysis system and set up to acquire 10X phase contrast images at a scanning interval of 60 minutes, scan type ‘Scratch Wound’, measuring relative wound confluence. Relative wound confluence was averaged between 12 replicates for each condition (mean + SD) and statistical significance was calculated by Students t-test.

### Melanoma Cohort – Transcriptomic data

Transcriptomic data were generated in 703 FFPE tumours from the Leeds Melanoma Cohort, using the Illumina DASL^®^ array.

## Acknowledgements

We gratefully acknowledge the participation of all patients and families in this study, and research coordination by Mrs Jane White. VAK, AT and the work presented in this study were funded by the Wellcome Trust (Grant WT104076MA). SP was funded by Caring Matters Now Charity and by the Newlife Foundation. The work was supported by the GOSHCC Livingstone Skin Research Centre, and by the UK National Institute for Health Research through the Biomedical Research Centre at Great Ormond St Hospital for Children NHS Foundation Trust, and the UCL GOS Institute of Child Health.

## Author contributions

SP designed and carried out cellular modelling, RPPA, RNAseq, qRT-PCR and Western blot validations, proliferation and migration assays and immunocytochemistry, analysis of all associated data, produced the figures, and wrote the online methods. LA-O and ACT carried out MLPA copy number validation, the pilot NGS panel, RT-PCR tissue arrays, and analysis of data. DL designed and advised on the cellular modelling, RPPA, RNAseq and validations. MH, JN and JNB contributed patients and samples, with phenotypic, genotypic, and expression data and the associated analysis, and reviewed the manuscript. LH and GE analysed the NGS panel copy number data. SH and AP analysed the RNAseq, RPPA and regional similarity search data and produced the relevant figures. LH and GE analysed the NGS panel copy number data. NW, HC and YX designed and carried out the PR70 phosphatase activity and pull-down experiments, and reviewed the manuscript. PA, JM, VMdS, CC, GT, JAPB, and SP contributed patients, samples and phenotypic information. SB, RW and SMB conducted some sequencing experiments. DB and DM advised and helped analyse immunocytochemistry. PB was involved in early discussions of the genetic data. CH contributed patients and samples. Michael H advised on proliferation and migration assays. LL contributed cell line expression data and analysis with the relevant figure. SL and DH contributed to analysis of the array CGH data and validation. JM, SS, MZ contributed control cohort data on copy number. MM, DZ and WLD helped optimise cellular modelling and validation. KP and VB contributed patients and samples and reviewed the manuscript. NS analysed immunohistochemistry. Paulina S performed knock-down experiments and analysis. PS was involved in early discussion of the genetic data and reviewed the manuscript. EH and GM contributed to early research design and to the manuscript. JD helped conceive and direct the cellular modelling and signalling pathway characterisation, and contributed to the manuscript. VAK conceived, designed and directed the research, designed and carried out the initial human genetics experiments, contributed patients, analysed data, and wrote the manuscript.

## Supplementary Materials

### Materials and Methods

#### Protein preparation, and holoenzyme assembly with PR70 and B’γ1 regulatory subunits via GST-mediated pull-down assay

Expression and purification of PP2A Aα (9-589), Cα (1-309), PR70 subunit, and Cdc6, and assembly of the PP2A core enzyme (Aα-Cα heterodimer) followed procedures described previously. Approximately 10 μg of GST-AC (core enzyme) was bound to 10μl of glutathione resin via GST tag. The resin was washed with 200μl assay buffer three times to remove the excess unbound protein. Then, the indicated amounts of PR70 and B’γ1 were added to the resin in a 200μl volume suspended in the assay buffer containing 25 mM Tris (pH 8.0), 150 mM NaCl, 1 mM CaCl2, and 3 mM DTT. The mixture was washed three times with the assay buffer. The proteins remained bound to resin were examined by SDS-PAGE, and visualized by Coomassie blue staining. All experiments were repeated three times. The level of binding was quantified using Image J, and results from three separate experiments were fitted in GraphPad Prism (GraphPad Software, Inc.).

#### Phosphatase assay

Purified GST-Cdc6 (49-90) was phosphorylated *in vitro* by Cyclin A/CDK2 (1/20 w/w) with 10 mM MgCl2 and 10x molar concentration of ATP for 1 hour at 30°C. The phosphorylated protein was purified by gel filtration chromatography (Superdex 200, GE Healthcare) to remove free ATP, followed by overnight cleavage of the GST-tag with TEV protease (1/20 w/w). The pCdc6 peptide was then separated from GST or uncleaved peptide using untrafiltration membrane (Millipore) with a 10 kDa cut-off. The phosphatase assay was performed in 100μl assay volume (rather than 50 μL) at room temperature for 15 minutes and stopped by the addition of 2x malachite green (50 μL) rather than 100 μL of 1x malachite green to increase the sensitivity of the assay. The phosphatase activity of 0.16 µM of PP2A core enzyme was measured using 2.2 µM pCdc6 peptide in a buffer containing 25 mM Tris pH 8.0, 150 mM NaCl, 3mM DTT, 50 µM MnCl2 and 1 mM CaCl2, in the presence and absence of the indicated concentrations of PR70 (108-519). The absorbance at 620 nm was measured after 10 min incubation at room temperature. The PR70 concentration-dependent PP2A phosphatase activity toward pCdc6 was fitted in GraphPad Prism (GraphPad Software, Inc.) by two separate simulations for two ranges of PR70 concentrations, lower versus higher than the molar concentration of PP2A core enzyme. The experiments were repeated three times, representative results were shown.

#### Mammalian cell culture, recombinant expression of PR70, and fractionation of cell lysate via gel filtration chromatography followed by western blot

293T and C6 glioma cells were cultured in Dulbecco’s modified Eagle’s medium (DMEM) supplemented with 10% fetal bovine serum (FBS), 100 units/ml of penicillin, and 100 μg/ml of streptomycin. The V5-tagged human PR70 (V5-PR70) was cloned into murine leukemia retroviral vectors harboring a CMV promoter for over-expression. Retroviruses harboring V5-PR70 were packaged in 293T cells, and were used to infect 293T or C6 glioma cells with 50-80% confluence for over-expression of V5-PR70. The infection efficiency of retroviruses was monitored by the fluorescence signals of RFP or GFP included in retroviral vectors. The level of recombinant V5-PR70 in holoenzyme versus as free subunit were determined by fractionation of cell lysate over gel filtration chromatography to examine co-migration of V5-PR70 with the scaffold A and PP2Ac subunits. The relative amounts of A, PP2Ac, and V5-PR70 in each fraction were determined by western blot using antibodies that specifically recognize A (Millipore), PP2A Cα subunit (Millipore, 1D6), and V5-tag (Millipore), followed by quantification of the western blot signals using Image J. The experiment was repeated three times; representative results are shown.

### Key Resources

**Figure S1.**
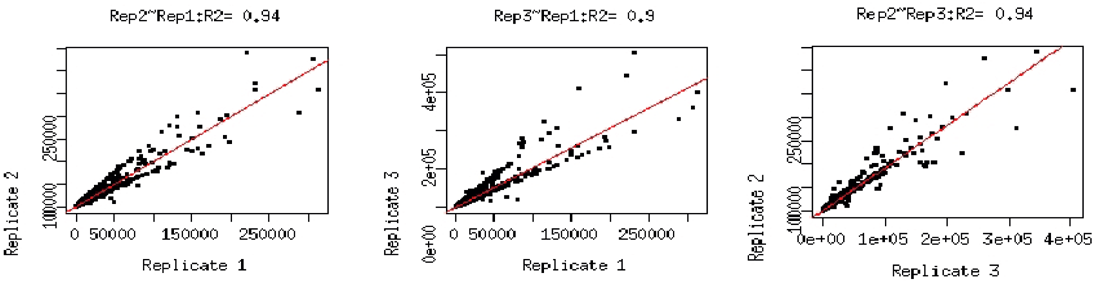
Replicates of induced and uninduced cells used in RNAseq show good correlation. From left to right, correlation between 1^st^ and 2^nd^ replicates, 1^st^ and 3^rd^ replicates and 2^nd^ and replicates.

**Figure S2.**
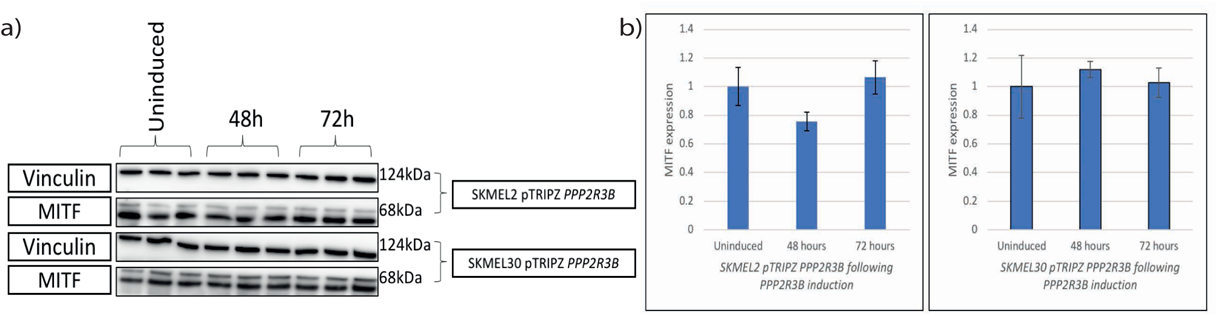
Overexpression of *PPP2R3B* does not induce proliferation via MITF. Western blot of uninduced and induced SKMEL2 pTRIPZ *PPP2R3B* and SKMEL30 pTRIPZ *PPP2R3B* cells for MITF expression **(a),** and quantification **(b),** show *PPP2R3B* overexpression does not consistently alter MITF activation across the cell lines in this system, suggesting the increased proliferation in cell lines observed following *PPP2R3B* overexpression is MITF independent

**Figure S3.**
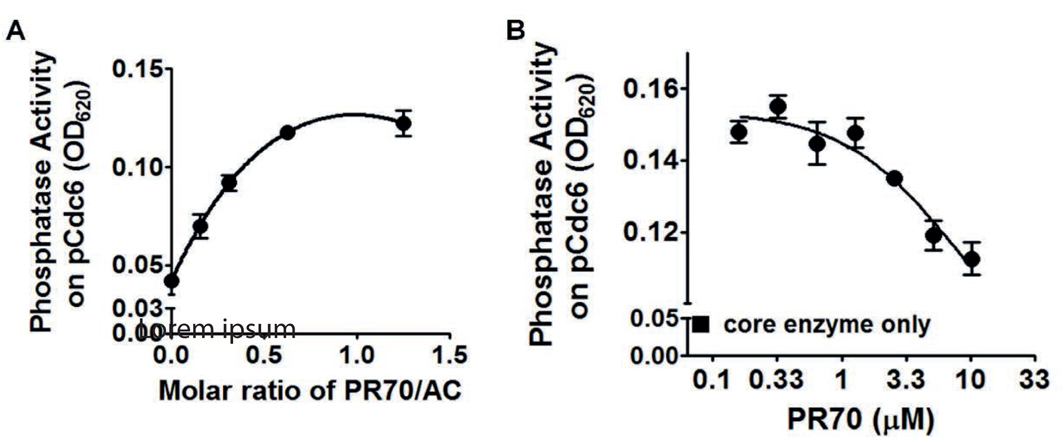
Increasing concentrations of PR70 gave increased PP2A activity toward its specific substrate, pCdc6, a cell cycle regulator that participates in tight control of replication licensing^41^ **(a)**. However, when the molar concentration of PR70 was higher than PP2A core enzyme, further increase of PR70 level reduced the phosphatase activity of PP2A, possibly due to a dominant negative effect via competitive inhibition **(b)**.

**Figure S4.**
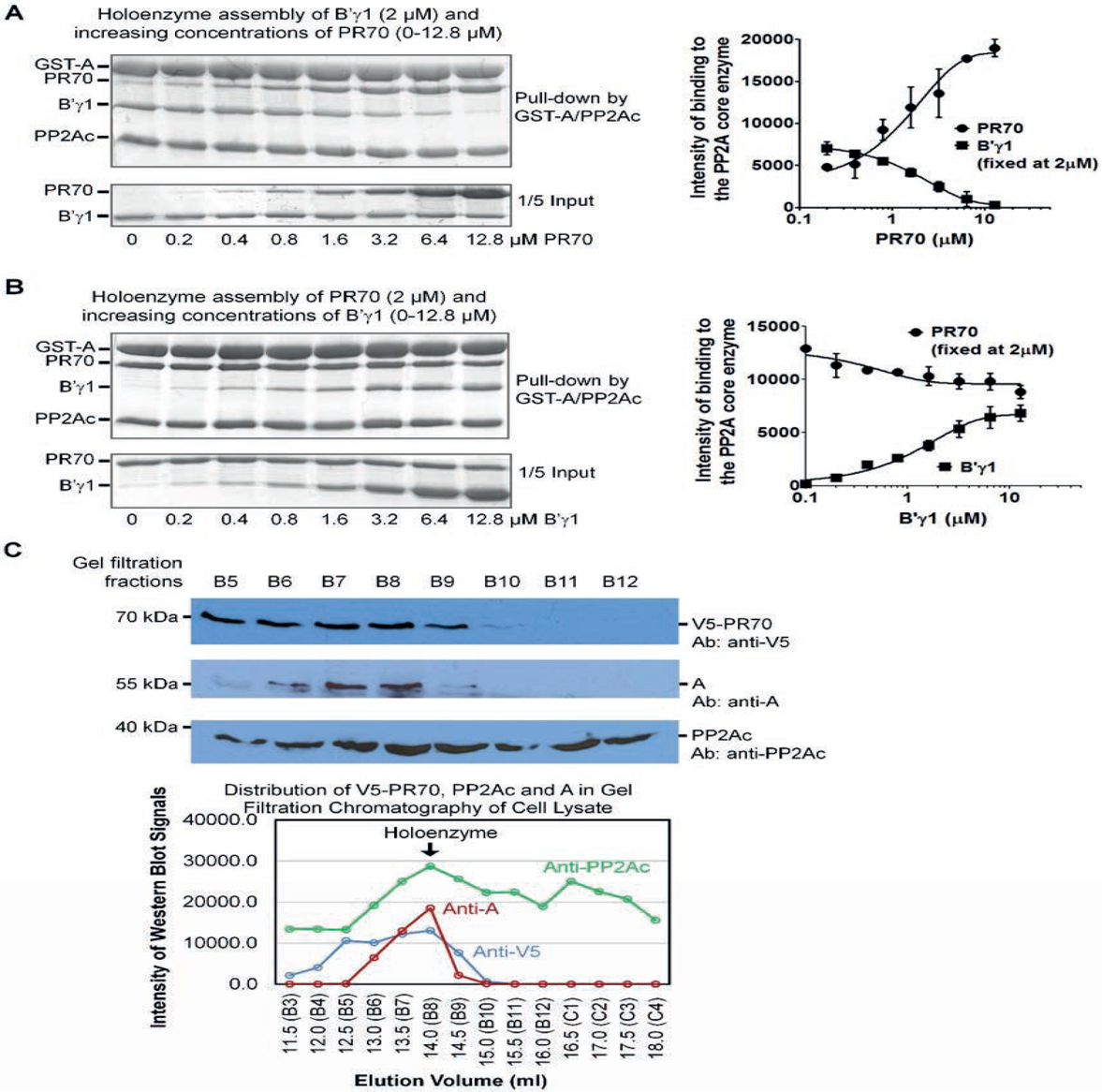
PR70 efficiently competes with the B’γ1 regulatory subunit for holoenzyme assembly. A). Pull-down of a fixed concentration of B’γ1 in the presence and absence of increasing concentrations of PR70 via GST-tagged PP2A core enzyme (A/PP2Ac). The bound proteins were visualized on SDS-PAGE by Coomassie blue staining. PR70 efficiently reduced the holoenzyme assembly with the B’γ1 regulatory subunit. B). Pull-down of a fixed concentration of PR70 in the presence and absence of increasing concentrations of B’γ1 similar to that described in A). B’γ1 could barely compete with PR70 for holoenzyme assembly. For both panel A) and B), the experiments were repeated three times; representative results are shown. The simulation of the quantified data from three repeats was shown on the right. C). Western blot detection of A, PP2Ac, and recombinant V5-PR70 in fractions of gel filtration chromatography of cell lysates with overexpressed V5-PR70. The distribution curves of the three components are shown in the lower panel.

**Table S1** – Upregulated genes associated with *PPP2R3B* expression are enriched for non-immune pathway signatures - transcriptomic data from 703 melanoma tumour samples from Leeds melanoma cohort (see methods for details)

**Table S2** – Down regulated genes associated with *PPP2R3B* expression are enriched for immune pathway signatures - transcriptomic data from 703 melanoma tumour samples from Leeds melanoma cohort (see methods for details)

**Table S3** – Pathway enrichment analysis from RNAseq of cell line inducible overexpression system for *PPP2R3B* (analysis excludes *PPP2R3B*), highlights suppression of the unfolded protein response and endoplasmic reticulum (ER) protein folding - comparison of pre- and post-induction samples in triplicate, at 6 and 16 hours post induction (see methods for details).

**Table S4** – Reverse Phase Protein Array (RPPA) raw data from cell line inducible overexpression system for *PPP2R3B* demonstrates enrichment for mammalian target of rapamycin (mTOR) and hypoxia-induced factor 1 (HIF-1) signalling pathways, with prominent biological signatures of response to heat and stress - comparison of pre- and post-induction samples in triplicate, at 6 and 16 hours post-induction (see methods for details).

**Table S5** – RNAseq data from cell line inducible overexpression system for *PPP2R3B*, significantly (Padj<0.05) differentially expressed genes in both cell lines in triplicate, at 6 and 16 hours post-induction (see methods for details). *C21orf91* was identified as the most significantly differentially expressed in both cell lines at both time points, other than *PPP2R3B*.

## Supplementary

### Melanoma Transcriptomic Data – Upregulated Pathways

The genes positively correlated with *PPP2R3B* are predominantly enriched in non-immune pathways, consistent with the lack of association between *PPP2R3B* expression and TILs (tumour infiltrating lymphocytes) or any specific immune cell score. There are only 24 pathways significantly enriched in these genes at FDR<0.05 (*Table S1*).

Using undirected interactions, the nodal genes in a network putting together all these pathways are: *MAPK11* (betweenness=2237) and *PLCB3* (betweenness=1120).

With directed interactions, the most important genes are *MAPK11* (betweenness=103) and *NFATC1* (betweenness=59).

**Table S1.** Melanoma Transcriptomic Data – Upregulated Pathways

| Rank | Pathway | FDR |
| --- | --- | --- |
| 1 | Rap1 signaling pathway(K) | 6.10E-04 |
| 2 | Axon guidance(K) | 6.10E-04 |
| 3 | ECM-receptor interaction(K) | 9.94E-04 |
| 4 | PI3K-Akt signaling pathway(K) | 3.49E-03 |
| 5 | DNA methylation(R) | 5.66E-03 |
| 6 | Pathways in cancer(K) | 9.73E-03 |
| 7 | Beta1 integrin cell surface interactions(N) | 0.0157 |
| 8 | Oxidative stress response(P) | 0.0209 |
| 9 | Arf1 pathway(N) | 0.0281 |
| 10 | Ras signaling pathway(K) | 0.0281 |
| 11 | Hedgehog signaling events mediated by Gli proteins(N) | 0.0281 |
| 12 | NoRC negatively regulates rRNA expression(R) | 0.0281 |
| 13 | EPH-Ephrin signaling(R) | 0.0281 |
| 14 | RNA Polymerase I Transcription(R) | 0.0308 |
| 15 | Signaling by VEGF(R) | 0.035 |
| 16 | MAPK signaling pathway(K) | 0.0388 |
| 17 | Focal adhesion(K) | 0.0388 |
| 18 | Endocytosis(K) | 0.0388 |
| 19 | Extracellular matrix organization(R) | 0.0388 |
| 20 | Shigellosis(K) | 0.0463 |
| 21 | EPHA forward signaling(N) | 0.0463 |
| 22 | DNA replication(P) | 0.0498 |
| 23 | Beta-catenin independent WNT signaling(R) | 0.0498 |
| 24 | Cell junction organization(R) | 0.0528 |
| 25 | Hedgehog signaling pathway(P) | 0.054 |

### Melanoma Transcriptomic Data – Downregulated Pathways

The genes negatively correlated with *PPP2R3B* expression are predominantly enriched in immune pathways. There are 80 pathways, top 25 of which are shown in Table S2.

Using undirected interactions, the top 5 nodal genes in a network putting together all these pathways are: *TGFB1* (betweenness=8395) and *AURKA* (betweenness=5133) and CD8A (betweenness=4893), *RAC2* (4880) and *LCK* (betweenness=4855).

With directed interactions, the most important genes are *LCK* (betweenness=174), *HLA-G* (betweenness=110), *MYD88* (betweenness=104), *TGFB1* (betweenness=86) and *CD8A* (betweenness=82).

**Table S2.** Melanoma Transcriptomic Data – Downregulated Pathways

| Rank | Pathway | FDR |
| --- | --- | --- |
| 1 | Natural killer cell mediated cytotoxicity(K) | 3.50E-08 |
| 2 | Antigen processing and presentation(K) | 2.33E-07 |
| 3 | Cytokine-cytokine receptor interaction(K) | 2.33E-07 |
| 4 | Immunoregulatory interactions between a Lymphoid and a non-Lymphoid cell(R) | 5.45E-07 |
| 5 | IL12-mediated signaling events(N) | 6.57E-07 |
| 6 | Malaria(K) | 6.57E-07 |
| 7 | Retinoid metabolism and transport(R) | 6.57E-07 |
| 8 | Graft-versus-host disease(K) | 1.17E-06 |
| 9 | IL12 signaling mediated by STAT4(N) | 1.28E-06 |
| 10 | Regulation of Insulin-like Growth Factor (IGF) transport and uptake by Insulin-like Growth Factor Binding Proteins (IGFBPs)(R) | 1.81E-06 |
| 11 | Cell adhesion molecules (CAMs)(K) | 3.05E-06 |
| 12 | Post-translational protein phosphorylation(R) | 1.77E-05 |
| 13 | Chemokine signaling pathway(K) | 3.29E-04 |
| 14 | G alpha (i) signalling events(R) | 3.29E-04 |
| 15 | Primary immunodeficiency(K) | 4.49E-04 |
| 16 | IL23-mediated signaling events(N) | 4.49E-04 |
| 17 | Osteoclast differentiation(K) | 6.46E-04 |
| 18 | Response to elevated platelet cytosolic Ca<sup>2+</sup>(R) | 8.66E-04 |
| 19 | Hematopoietic cell lineage(K) | 1.02E-03 |
| 20 | Th17 cell differentiation(K) | 2.12E-03 |
| 21 | TCR signaling in naive CD4+ T cells(N) | 2.12E-03 |
| 22 | African trypanosomiasis(K) | 2.32E-03 |
| 23 | TCR signaling in naive CD8+ T cells(N) | 3.26E-03 |
| 24 | Leishmaniasis(K) | 3.28E-03 |
| 25 | Extrinsic prothrombin activation pathway(B) | 3.54E-03 |

